# Wild common crossbills produce redder body feathers when their wings are clipped

**DOI:** 10.1101/2022.03.28.486028

**Authors:** Blanca Fernández-Eslava, Alejandro Cantarero, Daniel Alonso, Carlos Alonso-Alvarez

## Abstract

The animal signaling theory posits that conspicuous colorations exhibited by many animals have evolved as reliable signals of individual quality. Red carotenoid-based ornaments may depend on enzymatic transformations (oxidation) of dietary yellow carotenoids, which could occur in the inner mitochondrial membrane (IMM). Thus, carotenoid ketolation and cell respiration could share the same biochemical pathways. Accordingly, the level of trait expression (redness) would directly reveal the efficiency of individuals’ metabolism and hence, the bearer quality in an unfalsifiable way. Different avian studies have described that the flying effort may induce oxidative stress. A redox metabolism modified during the flight could thus influence the carotenoid conversion rate and, ultimately, animal coloration. Here, we aimed to infer the link between red carotenoid-based ornament expression and flight metabolism by increasing flying effort in wild male common crossbills *Loxia curvirostra* (Linnaeus). In this order, 295 adult males were captured during winter in an Iberian population. Half of the birds were experimentally handicapped through wing feather clipping to increase their flying effort, the other half being used as a control group. The rump feathers were also plucked in all the birds to stimulate plumage regrown of a small surface during a short time-lapse. Thirty-four birds with fully grown rump feathers could be recaptured in the subsequent weeks. Among them, male crossbills with experimentally impaired flying capacity showed redder rump feathers than controls. This result would support the hypothesis that the flying metabolism influences the redox enzymatic reactions required for converting yellow dietary carotenoids to red ketocarotenoids.

**Summary statement:** Carotenoid-based colorations may depend on pigment enzymatic transformations linked to mitochondrial function. The flight metabolism could influence this. Here, common crossbills with clipped wings produced redder feathers than controls.

## INTRODUCTION

Conspicuous colorations in many animal species have attracted the attention of evolutionary biologists from Charles Darwin. The current animal signaling theory posits that these traits may evolve as individual quality signals that act in sexual or social signaling contexts (Maynard Smith and Harper 2003). If evolving as individual quality signals, the traits should transmit reliable information favoring the receiver’s fitness (Maynard Smith and Harper 2003). It has been hypothesized that the signal reliability is maintained by the costs of trait production, preventing potential cheaters from correctly expressing the trait (i.e. handicap signals; Zahavi 1975, Grafen 1990; also e.g. a recent review in Penn and Számadó 2020). Additionally, or alternatively, signal reliability could be assured when a close link between the trait development process and the signaler’s intrinsic quality is established. These traits are known as index signals (Maynard Smith and Harper 2003). The most easily recognizable index signals are some beetles’ huge weapons (mandibles), big antlers of ungulates and frog or deer vocalization frequencies, which are all strongly correlated to individual body size (e.g. Bierniasky et al., 2014). All of them should allow inferring individual quality because body size determines individual fitness in these species (Maynard Smith and Harper 2003).

In the last years, another index signal type has been proposed. It is produced by pigments (red keto-carotenoids) whose synthesis could be produced at the core of the cell respiratory function, i.e. into the mitochondrion (Hill 2011; Johnson and Hill 2013; Cantarero and Alonso-Alvarez 2017). We should first mention that, in many avian species, the red colorations are generated by converting yellow carotenoids acquired with food into red keto-carotenoids (McGraw 2006; Johnson and Hill 2013). That transformation involves redox enzymatic reactions (McGraw 2006; Johnson and Hill 2013). The cited change is performed by a group of enzymes (ketolases) that seems to be located at the inner mitochondrial membrane (IMM; Johnson and Hill 2013; Hill et al. 2019). This means they would share the biochemical pathways involved in cell respiration (i.e. the shared-pathway hypothesis; see Hill 2011; Johnson and Hill 2013). Thus, the level of trait expression (redness) would directly reveal the basis of metabolic function and, therefore, the bearer quality in an unfalsifiable way (Hill 2011).

Otto Völker (1957) was probably the first to suggest that red carotenoid-based colorations are constrained by redox metabolism (see also Weber 1953 and 1961). He studied male common crossbills (*Loxia curvirostra*, Linnaeus), whose plumage color ranges from dull yellow to bright red, and feather redness is due to feather ketocarotenoid accumulation (Del Val et al. 2009a; Cantarero et al. 2020a). The researcher observed that the red males housed in cages could not replace their original red plumages and were always molted to produce yellow feathers. He suspected that, contrarily, crossbills allowed to fly could produce red feathers because they would have a better redox metabolism favoring the conversion of dietary yellow pigments (also Weber 1961).

Different avian studies have described that the flying effort may increase oxidative stress (e.g. Costantini et al. 2008; Jenni-Eiermann et al., 2014; Yap et al., 2017). Oxidative stress is usually defined as an imbalance between the production rate of reactive oxygen species (ROS) due to cell metabolism and the state of different antioxidant defenses (e.g. Sies and Jones 2007). It leads to oxidative damage, which is involved in senescence and age-related diseases (Barja 2014). An important part of ROS in cells is generated into the mitochondrion during cell respiration (St-Pierre et al., 2002; Zhang and Wong 2021). The flying effort, manipulated by training or inferred from migratory behavior, was positively correlated to mitochondrial oxidative stress in birds’ blood and pectoral muscles (i.e., Gerson et al., 2012; Gutierrez et al., 2019). It has also been correlated to higher superoxide dismutase 2 (a specific mitochondrial antioxidant enzyme) gene expression in avian pectoral muscles (Banerjee and Chaturvedi, 2016). Therefore, we can hypothesize that the redox metabolism modified during the flight could influence the carotenoid conversion rate and, ultimately, body coloration, such as early proposed by Völker (1957).

However, the evidence to the present date is weak and mostly based on a few avian studies where flying effort was increased and color expression measured. First, Schmidt-Wellenburg et al. (2008) augmented the flying workload of captive Rosy starlings (*Pastor roseus*; Linnaeus) by using a wind tunnel. Their pink plumage is generated by red ketocarotenoids (Galván et al. 2019), but no significant effect on plumage color after the natural molt was detected (Schmidt-Wellenburg et al. 2008). Leclaire et al. (2011) also increased the flying effort of wild adult kittiwakes (*Rissa trydactila*; Stephens) but in this case by clipping some wing feathers during reproduction. This technique has often been used to manipulate the cost of reproduction with different objectives (e.g. Slagsvold and Lifjeld 1990; Ardia and Clotfelter 2007; Cantarero et al., 2014). Leclaire et al. (2011) reported a brightness decline in the red bill gape of kittiwakes (a trait also colored by ketocarotenoids: see Leclaire et al 2019). However, they did not detect changes in the hue or saturation of the gape, which are color parameters linked to tissue carotenoid concentrations (McGraw 2006). Moreover, the red ketocarotenoids are obtained with food in the seabirds and directly deposited on ornaments without metabolic conversion (reviewed in McGraw 2006). Tarvin et al. (2016) performed a similar feather clipping in wild female American goldfinches (*Spinus tristis* Linnaeus) and did not detect bill color change when recaptured three weeks later. However, red ketocarotenoids have still not been described in the bill of this species. Lastly, a fourth study indeed detected a change in the color of a ketocarotenoid-based ornament. Wild male red-backed fairy-wrens (*Malurus melanocephalus* Latham) with clipped wings increased the surface of red plumage after natural molt compared to controls (Barron et al., 2013), which would agree with Völker (1957)’s hypothesis. However, the wing-clipped birds reported better body condition (inferred from a body fat score) than controls. The authors thus deduced that the red surface increment was due to lower mobility or higher food intake. The birds could not have endured a higher flying effort. Note that body mass loss is often detected in feather clipping studies (e.g. Lifjeld and Slagsvold 1988; Weimerkirsch et al., 1995; Lind and Jacobson 2001; Ardia and Clotfelter 2007; Harding et al 2009; Leclaire et al 2011; Matysiokova and Remes 2011; Wegmann et al., 2015; Casagrande and Hau 2018).

The present study aims to infer the link between red carotenoid-based ornament expression and flight metabolism by increasing flying effort in wild male common crossbills. In captive males of this species, we have recently described that the treatment with synthetic mito-targeted antioxidant (mitoTEMPO) favored the production of a redder plumage (Cantarero et al., 2020a). We have also found that wild male crossbills lose plumage redness with age, which may suggest a link to senescence and, accordingly, oxidative stress (Fernández-Eslava et al., 2021a). Redder crossbills also have higher probabilities of being recaptured (i.e., a proxy of survival; Fernández-Eslava et al., 2021a,b), thus linking plumage color to individual fitness. Importantly, we have recently found that redder males have proportionally longer flying feathers, thus probably correlating flying activity to carotenoid conversion (Fernández-Eslava et al., under review; see also Alonso and Arizaga, 2013).

We have used the same technique applied in other studies to increase flying effort (i.e. wing feather clipping). We also plucked the rump feathers in all the birds to stimulate plumage regrown in a small surface during a short time-lapse, a method previously used in this species (i.e. Cantarero et al., 2020a). We measured the rump color just before plucking it and when the birds were recaptured several weeks later. Almost 300 male common crossbills were captured throughout autumn-winter in the same Iberian population. This sample size was needed to obtain a minimum recapture sample allowing comparisons. We assume that our manipulation increased the flying effort. We should note that crossbills are social birds during the winter (e.g. Benkman 1997), clipped individuals being probably forced to increase their energy expenditure to follow the flock. In this regard, the individual body mass change was tested, a body mass loss being predicted in clipped birds but not in controls. In terms of plumage coloration, two alternative predictions can be formulated. First, a higher flying effort reduces plumage redness. This could be expected if wing-feather clipping reduces the capacity to acquire dietary yellow carotenoids or augments oxidative stress, decreasing carotenoid availability to feathers (e.g. carotenoids being bleached by ROS or consumed as antioxidants; e.g. Simons et al., 2012; Koch et al., 2018). A hypothetical flight-related oxidative stress could also directly impair ketolase activity, considering that high ROS levels alter the mitochondrial metabolism (reviewed in e.g. Barja 2014). Alternatively, the manipulation could increase plumage redness. If joined with a body mass loss, this outcome would agree with a scenario where exercise raises redox rates (often described in mammals; e.g. Campos et al., 2013; Park et al., 2016), favoring carotenoid conversion, such as early envisaged by Völker (1957).

## MATERIALS AND METHODS

### Experimental design

The manipulation of free-ranging male crossbills required many captures, allowing to recapture enough experimental and control birds. This was only feasible during autumn and winter when birds group in flocks and crossbill captures rise. In that period, the birds are mostly resident individuals with high recapture probabilities (Alonso and Arizaga 2011, 2017). According to this, the manipulation period was performed from October 19, 2019, to February 14, 2020. This allowed capturing 295 adult male crossbills in the same ringing station (El Royo, Soria, Spain) throughout 17 sampling sessions. All the birds were captured in two 12 long 16 mm mesh mist-nets placed close to sites traditionally used to provide salt for livestock (Fernandez-Eslava et al. 2020).

The captured birds were ringed, and their sex and age EURING code were notated (based on Jenni and Winkler, 1994 and Svensson, 1998). The minimum age in months was subsequently established based on plumage features following Fernández-Eslava et al. (2021a). The birds were then weighed (±0.1 g) and several biometric variables were measured with a digital caliper. These included the wing length (± 0.5 mm, method III of Svensson, 1998), tail length (± 0.5 mm), tarsus length (± 0.01 mm) and head length (± 0.01 mm). The subcutaneous fat extension and pectoral muscle shape were registered in a visual score (fat: 0-8, with 0.25 subclasses in Kaiser 1993; muscle: 0-3 scale, in Bairlein 1995) and always by the same person (DA). Lastly, a four-level score was used to classify each bird’s plumage color, and two digital pictures of each individual were taken (see *Color measurements* below).

Birds were then assigned to the control or feather-clipped group. In the latter group, primaries 3 and 5 of each wing (counting inward from the distal wing margin) were clipped at the base of the rachis with scissors in feather-clipped males. Control males were similarly handled but without feather clipping. Additionally, the rump feathers of both groups were plucked after taking pictures (see below) to stimulate plumage regrown. The group assignation was *a priori* made randomly. However, we also attempted to balance the treatments regarding the cited plumage color categories and the bird’s age. Thus, finally, 151 controls and 144 clipped-feather birds were assigned. Only three controls and one clipped bird were ending their wing molt at the time of manipulation (treatment x molt occurrence: *χ*^2^ = 0.92, df = 1, *p* = 0.337). Among these four birds, only one control bird was recaptured. The results were similar when excluding these four individuals from the analyses.

Recaptures took place from January 2 to July 24, 2020. This means that some birds could be recaptured during the period dedicated to treatment assignment and manipulation of other birds. Only those recaptures that took place more than three weeks after the first capture were included in the statistical analyses. This was the minimum period observed to allow birds to regrow their rump feathers fully. Five recaptures produced within less than three weeks were accordingly discarded as the rump was not regrown (lapse mean ± S.D: 81 ± 9.3 d, range 21-208 d). The first birds showing a natural molt of the rump were found after July 24. Thus, only the color of the feathers induced by the experimental plucking was analyzed—a full inspection of body plumage from pictures and wing feather molting patterns allowed to verify this. Thirty-four recaptured birds met these criteria (20 control and 14 feather-clipped birds). The probability of being recaptured did not differ between the two groups tested in the analyses (*χ*^2^ = 0.89, df = 1, *p* = 0.344).

All the experimental work was approved by the government office in charge of the ethics in animal experiments (i.e., Organo Competente Consejeria de Medio Ambiente de la Comunidad de Madrid; approval ref. PROEX 103.4/20).

### Color measurements

Digital photographs of the breast and the back of red crossbills were always taken by putting the birds at the same position and fixed distance from the objective (Canon Macro Lens EF 50 mm; see also Cantarero et al., 2020a; see Fig. 1). A Kaiser Repro Base (Kaiser Fototechnik, Buchen) was used, including a gridded board and a column to place the camera at the same height (distance from the board to the lens: 38 cm). The base was covered with opaque grey cellular polycarbonate sheets placed vertically to cover the four sides of the board. It allowed us to enter the camera objective and ring flash from the top (Canon Macro Ring Lite MR-14EX). The sheets were perforated to allow entering the hands of a person that would hold the bird’s body resting on the board surface. All the open surfaces were covered with PVC blackout fabric to reduce the light on the board surface. The focus and diaphragm of the camera and the ring flash were all manually fixed to avoid the interference of automatic functions.

**Figure 1.**
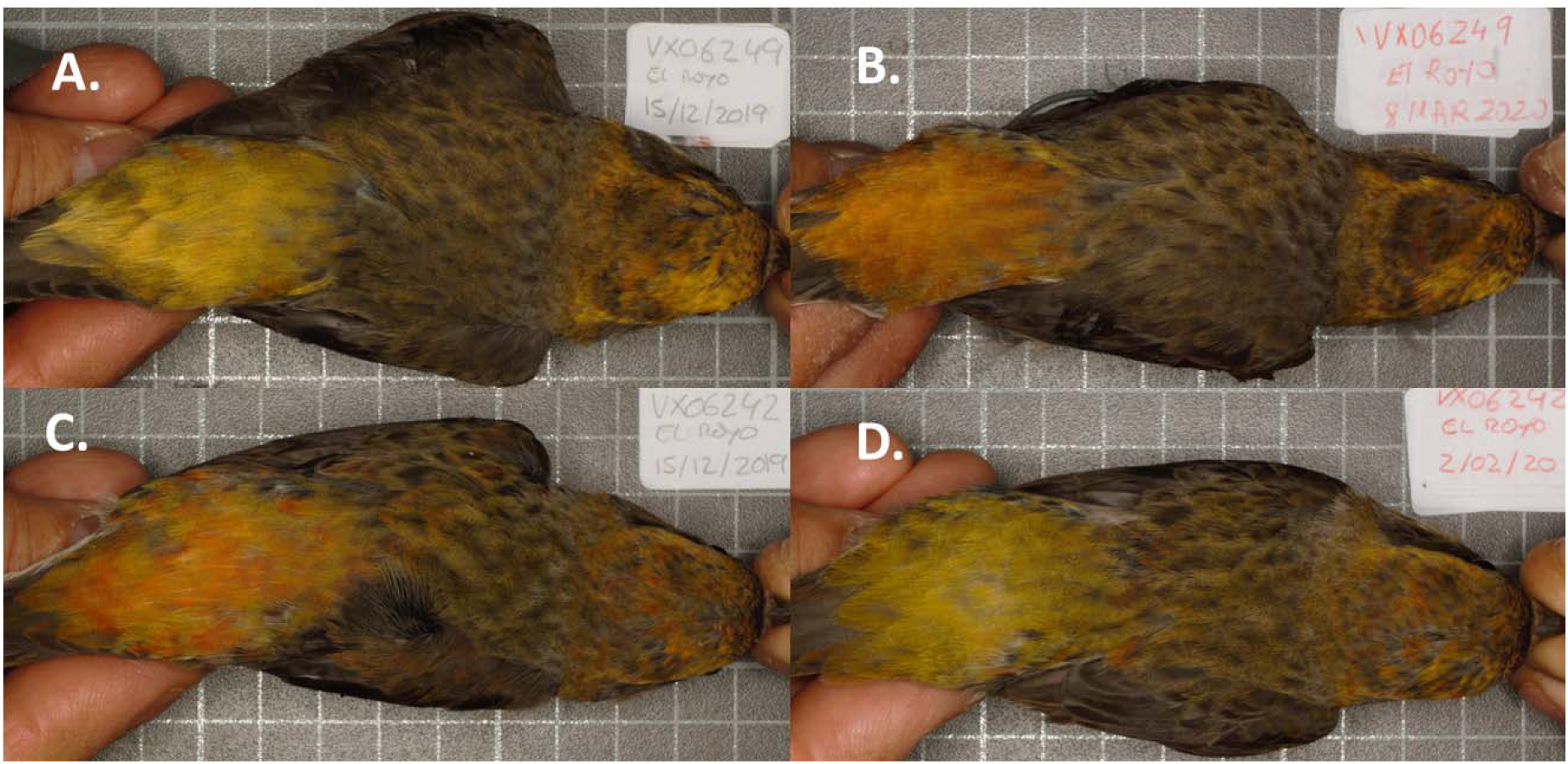
Example of plumage color change. Plumage color variation between capture and recapture dates in a clipped bird (A., B.) and a control individual (C., D.) before and after manipulation (from the left to the right). The new rumps always ranged from yellow to orange colors (see also **Figure 3**).

Two digital pictures of each bird were taken (i.e. one including the breast and abdomen, and another the backside). Thus, the bird was placed in face-up or prone positions by pulling the bill and legs. A standard grey card (Kodak, NY, USA) was used as a reference, placed next to the bird’s flank on the board surface, and always in the same position. Digital photographs were standardized and analyzed using ‘SpotEgg’ software (Gómez and Liñán-Cembrano 2017), which allows the user to manually draw any region and provide information about its coloration, size or shape. Areas with disorganized feathers were avoided. For each individual, the average of red, green, and blue components of the colored area of the chest (1), the head and back (2), and rump (3) were separately calculated. The rump area in the picture of the first capture was similar to that measured on the new rump and highly correlated (Pearson’s *r* = 0.83, *p* <0.001). We then determined each area’s hue (º), chroma and brightness through the Foley and van Dam (1982) algorithms. In a previous study on the same species, the repeatability (Lessells and Boag 1987) of these three variables taken twice was very high (all *r* > 0.90, *n* = 30; Cantarero et al., 2020a). Finally, since a low hue indicates a redder color, we reversed their values (i.e. multiplying by -1) to obtain a more intuitive variable, i.e. “redness.” We rescaled the variable by adding the minimum negative value to obtain positive data. Thus, high plumage redness values indicate redder traits (e.g. Cantarero et al., 2020a,b). The rump hue measures taken from another sample of crossbills reported a high correlation to red ketocarotenoid concentrations in rump feathers (Supplementary Information).

In addition, the birds were also classified in one of the four-color categories described by del Val et al. (2009b), i.e., yellow, patchy, orange or red. These categories consider the whole body area (Fernández-Eslava et al., 2021a). This body color score allowed to balance the treatment assignation in terms of individual coloration. The score correlates with objective colorimeter measures (del Val et al. 2009b) and is repeatable when assessed twice by the same person (DA) naïve to bird identity (measures separated by one week; see Cantarero et al. 2020a). Moreover, the score also positively correlated with the concentration of red ketocarotenoids in feathers, but not with yellow carotenoid levels (Supplementary Information).

### Statistical analyses

The analyses were performed with SPSS version 27. In the initial capture (i.e. *N* = 295), the sampling date, minimum age (i.e. Fernández-Eslava et al., 2021a), and wing, tail, and bill lengths, did not follow a Gaussian distribution (Shapiro-Wilk tests), and arithmetic procedures could not normalize them. Accordingly, treatment comparisons were made by non-parametric Mann-Whitney’s U tests in these variables. The body mass and tarsus length were normally distributed and comparisons were tested by Student’s t-tests. An ANCOVA model testing initial body mass as dependent, the treatment as a fixed factor, and morphometries as covariates was also performed to test potential biases in size-corrected body mass at the first capture (only covariate terms at *p* < 0.10 were retained in the model after a backward stepwise procedure).

In the dataset only including the recaptured birds (i.e. *n* = 34), the initial date of capture, number of days elapsed to the recapture and tail length were not normally distributed and compared by Mann-Whitney’s U tests. The initial body mass, body mass change to recapture (%) and other morphometries followed a Gaussian distribution and were analyzed by Student’s t-tests. Size-corrected body mass was tested using an ANCOVA, testing body mass as the dependent variable, the treatment as a fixed factor and significant body morphometries as covariates. Also, in this subset, the hue, chroma or brightness at the first or second capture dates, and in any body part, were normally distributed. However, the hue of the rump did not follow a normal distribution, and it could not be normalized by arithmetic transformations, being tested by Mann-Whitney’s U or Wilcoxon matched-pairs tests. Bivariate Spearman’s correlation coefficients were used when at least one of the variables was not normal. In Student’s *t*-tests, homoscedasticity was analyzed by Levene’s *F* tests. Two-tailed tests were always used.

## RESULTS

### Testing potential initial biases

Among the initially captured birds, the minimum age in months, initial date of capture and plumage color categories were not significantly biased between treatments (Mann-Whitney’s *U* = 10232, *p* = 0.379, *U* = 9804, *p* = 0.143 and *χ*^2^ = 4.78, df = 4, *p* = 0.310, respectively). Moreover, the wing, tail and bill lengths did not differ (all MW tests *p* > 0.240) and neither the body mass or tarsus length (*t* = 0.939, df = 286, *p* = 0.349 and *t* = 0.872, df = 292, *p* = 0.384, respectively). Size-corrected body mass was also balanced between treatments at the first capture (treatment: *F*_1,278_ = 0.01, *p* = 0.978; tarsus length: *F*_1,278_ = 19.1, *p* < 0.001; wing length: *F*_1,278_ = 13.8, *p* < 0.001). Finally, the muscle and fat scores did not differ (Mann-Whitney’s *U*, both *p* > 0.80)

In the subsample of recaptured birds (i.e. *n* = 34), no significant difference in rump hue (redness), saturation or brightness was found at the first capture (*U* = 119, *p* = 0.462, *t* = -0.93, df = 32, *p* = 0.361 and *t* = 0.17, df = 32, *p* = 0.863, respectively). Similarly, no color component of any other body region (i.e. head and back or breast areas) reported any significant difference between treatments (all *t*-tests: *p* > 0.13). Moreover, the minimum age in months, the initial capture date, the number of days elapsed from that date to the recapture, and body morphometries including size-corrected body mass did not significantly differ with treatments in this subset (all tests *p* > 0.20). Only the initial body mass showed a trend to statistical significance (*t* = 1.75, df = 32, *p* = 0.090), with clipped birds tending to be heavier than controls (means± SD: 38.6 ± 1.5 g and 37.4 ± 2.2 g, respectively). However, initial body mass did not correlate with rump redness at the first capture or recapture (Spearman’s *r* = - 0.202, *p* = 0.251 and *r* = 0.08, *p* = 0.654, respectively). Lastly, muscle and fat scores did not differ (Mann-Whitney’s *U*, both *p* > 0.74)

### Experimental effects

The body mass change (%) from the initial to the second capture clearly differed between control and clipped birds (i.e. *t* =2.97, df = 32, *p* = 0.006; Cohen’s *d* effect size: 1.04; see Fig. 2). Clipped birds lost about 1.7% of their initial body mass (difference to zero: *t* = -3.03, df = 13, *p* = 0.010), whereas controls showed a statistical trend to increase body mass in a similar proportion (*t* = 1.96, df = 19, *p* = 0.065; Fig. 2). The muscle score at recapture ranged from 1 to 3 in the 0-3 scale, and did not differ between treatments: U = 91, *p* = 0.236). The subcutaneous fat score did not show variability, being untested (always a 0.5-value, small stripe into the furcular depression).

**Figure 2.**
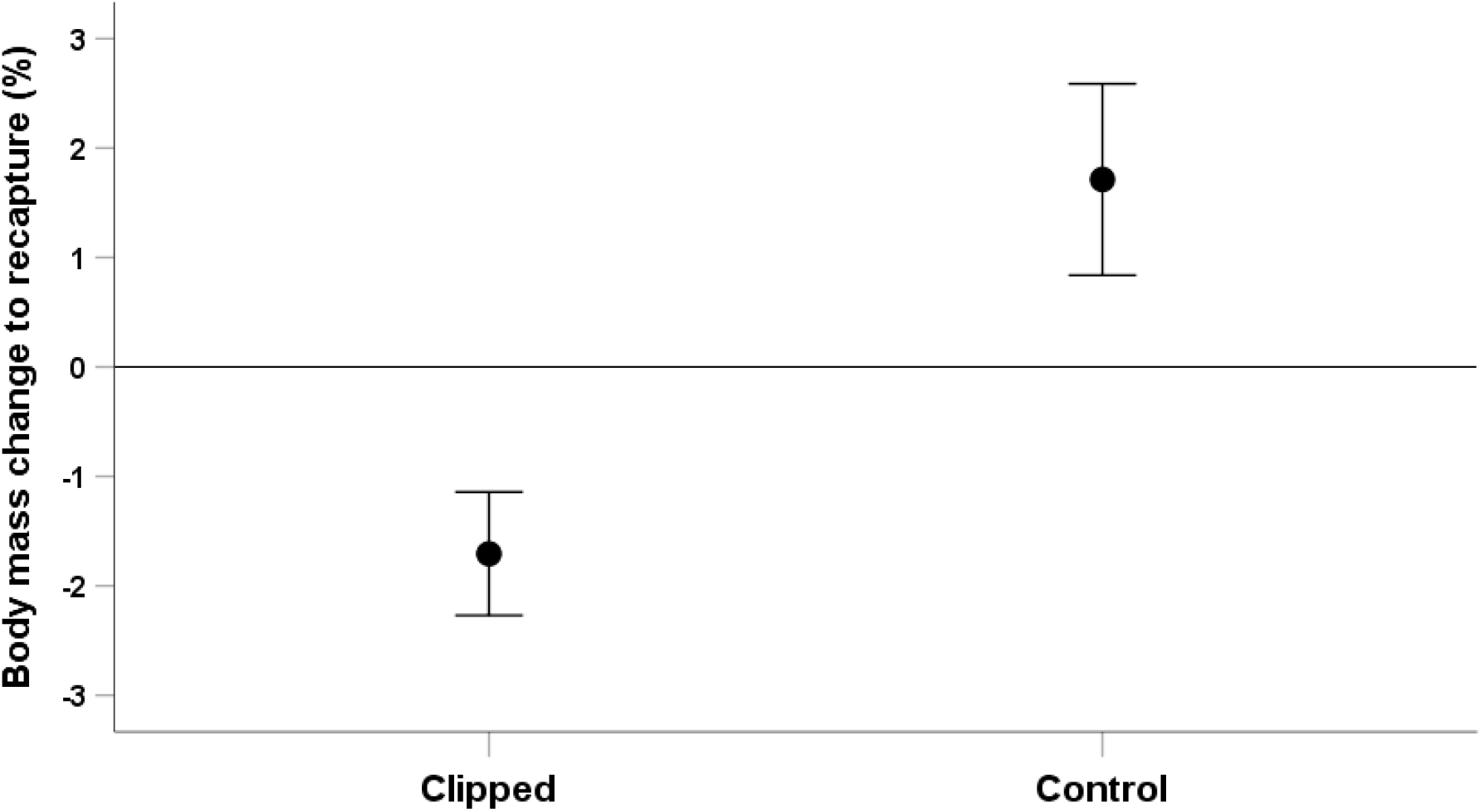
Reduced body mass in wing-clipped crossbills. Body mass change (%) from the initial capture to recapture in adult male common crossbills used as controls or being manipulated to increase their flying effort by clipping two primary feathers (i.e. third and fifth) of each wing (see Methods; means ± S.E.M.).

The redness of the rump strongly declined from the first to the second capture (Fig. 3; *Z* = -3.92, *p* < 0.001 and *Z*= -2.79, *p* < 0.001; Wilcoxon matched-pairs tests for control and clipped birds, respectively). The redness of the regrown rump redness was uncorrelated to the original value (Spearman’s *r* = 0.126, *p* = 0.477). This could be due to reduced redness variability (SD dropped from 5.95 to 2.98) and the experimental effect (Fig. 3). Thus, a positive correlation seems to arise among controls but not in clipped birds (*r* = 0.395, *p* = 0.085 and *r* = -0.125, *p* = 0.669, respectively). Despite the evident color decline and lack of within-individual correlation, the redness of the new regrown feathers significantly differed between treatments (*U* = 80.5, *p* = 0.037), with clipped birds developing a redder rump than controls (Fig. 3).

**Figure 3.**
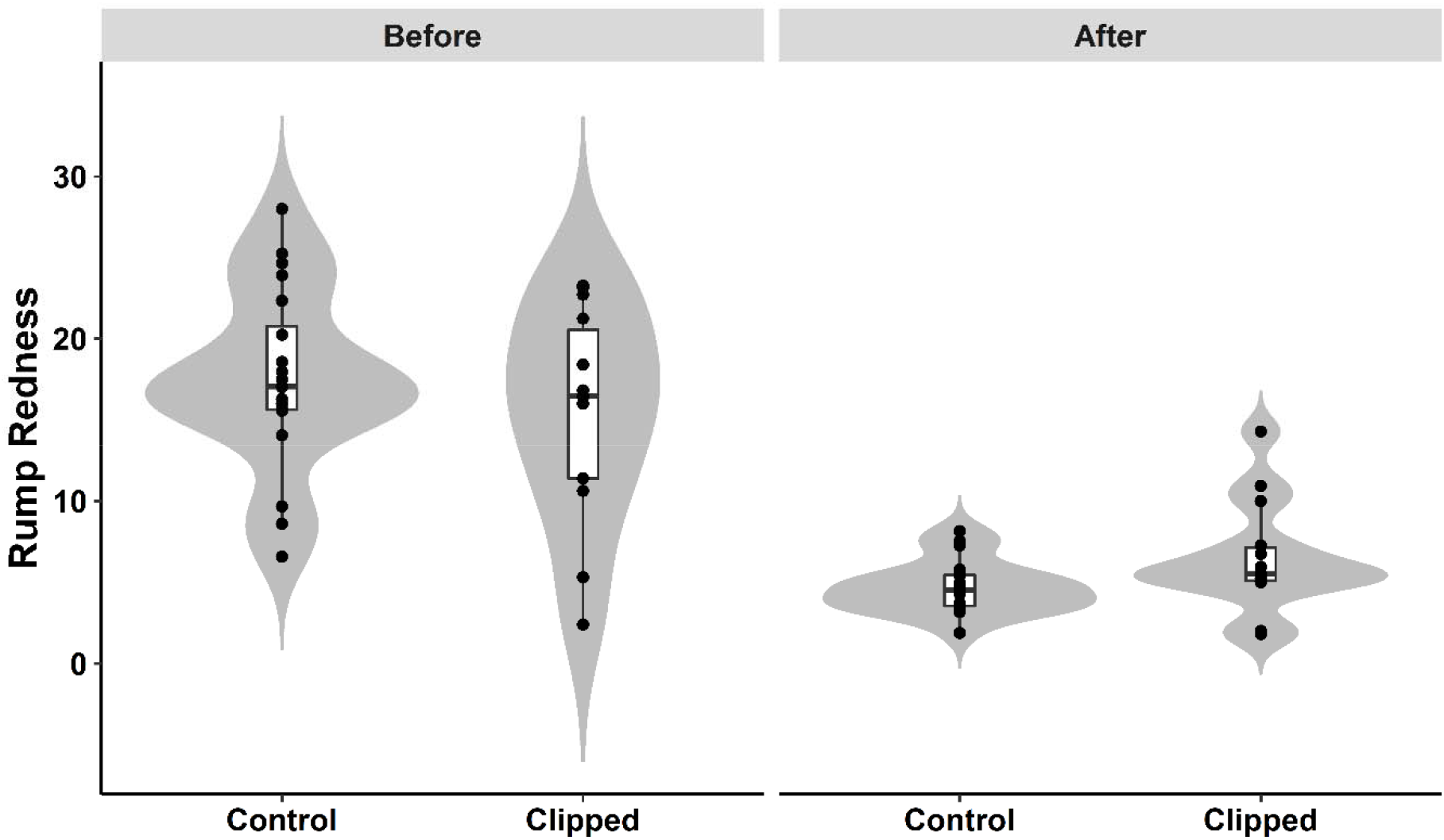
Redder rump feathers in wing-clipped crossbills. Rump color of wild male crossbills before (left) and after (right) wing feather clipping. Raw values (dots), medians, quartiles, and ranges are shown. The violin plot represents the density function. (control: *n* = 20; clipped: *n* = 14).

## DISCUSSION

Here, male crossbills with experimentally impaired flying capacity and, probably, obligated to alter their metabolism (Fig. 1) produced redder feathers than controls (Fig. 2 and 3). The red coloration of male common crossbills is due to the accumulation of red ketocarotenoids (mainly 3-hydroxyechinenone) in feathers, which requires enzymatic oxidation (ketolation) of yellow dietary carotenoids (del Val et al., 2009a; Cantarero et al., 2020a). Thus, the main result mostly supports the hypothesis that flying effort influences carotenoid conversion.

Control male crossbills tended to gain weight during the study (Fig. 1). Increased body mass and fat reserves from winter to spring have been documented in captive and wild common crossbills from North America (Hahn 1995; Cornelius and Hahn 2012), but this effect was non-significant in North Spanish populations (i.e. Alonso and Arizaga 2011). In contrast to controls, clipped crossbills lost body mass. A body mass loss of males wing clipped during the breeding period has been reported in many species and mostly interpreted as a consequence of increased flying effort due to reduced lift (e.g. Lifjeld and Slagsvold 1988; Weimerkirsch et al., 1995; Harding et al., 2009; Leclaire et al 2011; Wegmann et al., 2015; Casagrande and Hau 2018). Reduced body mass was also detected in captive wing-clipped tree sparrows (*Passer montanus* Linnaeus) during their molt, pectoral muscles increasing mass probably due to higher flight activity (Lind and Jacobson 2001). Here, fat and muscle scores did not provide any clue, but they reported low variability and were probably insensitive to subtle changes.

Alternatively, the clipped birds could have reduced body mass to compensate for the high flying cost imposed by increased wing loading. Compensatory body mass loss has mostly been described during the nestling-feeding period and interpreted as a way to optimize foraging efficiency (e.g. Moreno 1989; Merkle and Barclay 1996; Slagsvold and Johansen 1998; but see also Lind and Jacobson 2001). Crossbills have the main breeding peak between February and March (Alonso and Arizaga 2011). Unfortunately, we did not know the reproductive status of each bird. Nonetheless, at first capture, a rough index of active reproduction (i.e. cloacal protuberance occurrence; Schut et al., 2012) was registered. Among the recaptured birds, it showed high incidence (28/34: 82%) and equal distribution between both treatments (*χ*^2^ = 0.23, df = 1, *p* = 0.628). Therefore, we cannot infer that clipped individuals were engaged in different body mass regulation, although it cannot be fully discarded. However, we should note that a programmed body mass loss should involve reduced food intake (i.e. programmed anorexia; Jones 1994), probably constraining carotenoid ingestion and limiting pigment conversion. Thus, in our view, the most parsimonious interpretation of reduced body mass in clipped birds is that it was indeed a consequence of higher energy consumption to face flying effort.

Interestingly, the redness value of the new rump was approximately three times lower than that found at the first capture (Fig. 2). The birds were only able to regrown yellowish to orange feathers. We propose three non-excluding explanations. First, birds were mostly captured out of the molting season (see Alonso and Arizaga 2011; Fernández-Eslava et al., 2019). The natural molt involves a specific endocrine profile in birds (reviewed in Dawson 2015), and hormones could have constrained carotenoid conversion. For instance, high thyroid hormone levels are involved in avian molt activation (Gulde et al. 2009; Dawson 2015; Pérez et al., 2018). In captive male zebra finches (*Taeniopygia guttata* Vieillot), a triiodothyronine (T3) treatment triggered the molt of body feathers (Cantarero et al., 2020b). Simultaneously, the bill redness of these birds (a red-ketocarotenoid-based trait; McGraw and Toomey 2010) was affected by the interaction of a mitochondria-targeted antioxidant (mitoTEMPO) treatment and T3 dosage (Cantarero et al., 2020b). Thus, although the carotenoid-based ornament of zebra finches is not a plumage trait, the result suggests that the mitochondrial function could require a certain endocrine profile to favor pigment conversion. Second, low autumn-winter temperatures could affect redox metabolism. Low winter temperatures increase oxygen consumption and heart rates in common crossbills (Dawson and Tordoff 1964; Draud 2019). A different metabolism could then force a trade-off favoring survival over color expression, perhaps biasing mitochondrial metabolism from cell respiration linked to carotenoid conversion to heat production. Nonetheless, such a mitochondrial uncoupling is still unclear in birds (e.g. Walter and Seebacher 2009; Teulier 2010; Salin et al., 2015). Third, a low availability of dietary substrate carotenoids during winter could have limited carotenoid transformation. In male American redstart (*Setophaga ruticilla* Linnaeus), a hypothetical scarcity of carotenoids in food was also argued when interpreting decreased red chroma detected in a tail feather plucked and regrown during winter (Tonra et el. 2014). Moreover, in another close-related crossbill (i.e. *Loxia leucoptera* Gmelin), circulating β-cryptoxanthin levels (substrate for red ketocarotenoids in common crossbills; del Val et al., 2009a; Cantarero et al., 2020a) decrease out of the molting months, suggesting a decline in the availability of dietary sources of pigments (Deviche et al., 2008).

Whatever the factors constraining carotenoid conversion during the study, the new regrown plumage produced enough color variability to detect a significant experimental effect (Fig. 2). Contrarily to the prediction that increased flying effort could constrain carotenoid acquisition or impair redox transformations due to higher oxidative damage, leading to less red plumage, clipped crossbills produced redder (or more orange) feathers than controls. We highlight that, in a similarly short color range (yellow to orange) derived from a plucking experiment under captivity (Cantarero et al., 2020a), a positive correlation (Spearman’s *r* = 0.53) between crossbill rump redness and feather red ketocarotenoid concentration was detected (also Supplementary information). This suggests that the effect found here was indeed due to a higher activity of oxidase enzymes in charge of carotenoid biosynthesis (i.e. ketolases; e.g. Lopes et al., 2016) among wing-clipped birds. Thus, the result seems to support a link between flight-related metabolism and carotenoid conversion. At first glance, it coincides with Barron et al. (2013)’s study where wing-clipped male red-backed fairy-wrens produced larger red plumage areas than controls (the red color is also due to ketocarotenoids in this species; Rowe and McGraw 2008). Their analyses focused on recaptured birds (ten controls and ten clipped birds), surprisingly finding that only control birds lost body fat during the study (Barron et al., 2013). The authors attributed this to reduced activity or more time to acquire carotenoids, but not to higher flight effort. A link to flying metabolism was, thus, not considered. Moreover, the study did not report the number or characteristics of all the wrens initially manipulated or used as controls, potential biases or differential recovery rates being untested.

In the context of animal signaling theory, our result may suggest that common crossbill coloration could inform signal receptors about the intrinsic quality of the bearer in terms of cell metabolism (i.e. Hill 2011; Hill et al., 2019). However, it could also reveal a general phenotypic profile, including the flying capacity and other correlated traits linked to fitness. In this line, Chui et al., (2011) found that male golden-crowned kinglets (*Regulus satrapa* Lichtenstein) with redder crowns colored by ketocarotenoids left for migration earlier than yellow-crowned individuals, probably allowing early reproduction in breeding grounds and better reproductive success (e.g. Potti 1998). In the same line, Mateos-Gonzalez et al., (2014) found that redder male house finches (*Haemorhus mexicanus* Müller) were better able to escape from capture into the aviaries than yellowish birds, suggesting that red birds are better at avoiding predation. Note that the house finch plumage is colored by the same red transformed ketocarotenoid (3-hydroxy-echinenone) that male common crossbills deposit on their plumage (McGraw 2006; Del Val et al., 2009a; Cantarero et al., 2020a). Moreover, we have recently found that redder male crossbills have proportionally longer flight feathers than yellow birds (Fernández-Eslava et al., under review; see also Alonso and Arizaga, 2013) and more probabilities of being recaptured in the wild (an index of survival and, hence, of individual quality; Fernández-Eslava et al., 2021a,b).

In summary, the flying effort seems to favor carotenoid conversion and the production of redder plumage in male common crossbills. This is an experimental approach forcing all the clipped birds to afford the challenge during a relatively short time-lapse (a few weeks). Natural covariation between carotenoid-conversion rates, coloration and flying capacity could have evolved so that the ornament might reveal a general phenotypic profile including a certain flight-related metabolism. To confirm this, more correlational and experimental studies in free-ranging birds captured must be performed. Moreover, future work in this or similar animal models should specifically focus on cell respiration metabolism and ketolase activity rates during the trait development.

## Acknowledgements

We acknowledge the personnel of the Autonomous region government Comunidad de Castilla Leon (Spain) for authorizing the captures in the Sorian crossbill population.

## Competing interest

No competing interest is declared.

## Funding

The study was partially funded by Ministerio de Ciencia e Innovación (project ref. PID2019-109303GBI00, MCIN/AEI/10.13039/501100011033).

## Data availability

Data will be available at DIGITAL.CSIC public repository upon acceptance (DOI pending).

## Notes

### Competing Interest Statement

The authors have declared no competing interest.

## References

Alonso, D. and Arizaga, J. (2011). Seasonal patterns of breeding, moulting, and body mass variation in Pyrenean common crossbills Loxia curvirostra curvirostra. Ringing and Migration 26, 64–70. doi:10.1080/03078698.2011.587253

Alonso, D. and Arizaga, J. (2013). The impact of vagrants on apparent survival estimation in a population of common crossbills (Loxia curvirostra). Journal of Ornithology 154, 209–217. doi:10.1007/s10336-012-0887-2

Alonso, D. and Arizaga, J. (2017). Seasonal abundance patterns of common cross-bills Loxia curvirostra L., 1756 in two localities of the Navarran Pyrenees and implications for its survey through ringing. Munibe 65, 95–105. doi:10.21630/mcn.2017.65.08

Ardia, D. R. and Clotfelter, E. D. (2007). Individual quality and age affect responses to an energetic constraint in a cavity-nesting bird. Behavioral Ecology 18, 259–266. doi:10.1093/beheco/arl078

Bairlein, F. (1995) Manual of field methods, European-African songbird migration network. Wilhelmshaven, Germany: Institut für Vogelforschung.

Banerjee, S. and Chaturvedi, C. M. (2016). Migratory preparation associated alterations in pectoralis muscle biochemistry and proteome in Palearctic-Indian emberizid migratory finch, red-headed bunting, Emberiza bruniceps. Comparative Biochemistry and Physiology D. 17, 9–25. doi:10.1016/j.cbd.2015.11.001

Barja, G. (2014). The mitochondrial free radical theory of aging. Progress in Molecular Biology and Translational Science (1st ed., Vol. 127). Elsevier Inc. doi:10.1016/B978-0-12-394625-6.00001-5

Barron, D. G., Webster, M. S. and Schwabl, H. (2013). Body condition influences sexual signal expression independent of circulating androgens in male redbacked fairy-wrens. General and Comparative Endocrinology, 183, 38–43. doi:10.1016/j.ygcen.2012.12.005

Biernaskie, J. M., Grafen, A. and Perry, J. C. (2014). The evolution of index signals to avoid the cost of dishonesty. Proceedings of the Royal Society B 281, 20140876.

Campos, J. C., Gomes, K. M. S. and Ferreira, J. C. B. (2013). Impact of exercise training on redox signaling in cardiovascular diseases. Food and Chemical Toxicology, 62, 107–119. doi:10.1016/j.fct.2013.08.035

Cantarero, A. and Alonso-Alvarez, C. (2017). Mitochondria-targeted molecules determine the redness of the zebra finch bill. Biology Letters, 13(10). doi:10.1098/rsbl.2017.0455

Cantarero, A., López-Arrabé, J., Palma, A., Redondo, A. J. and Moreno, J. (2014). Males respond to female begging signals of need: A handicapping experiment in the pied flycatcher, Ficedula hypoleuca. Animal Behaviour 94, 167–173. doi:10.1016/j.anbehav.2014.05.002

Cantarero, A., Mateo, R., Camarero, P.R., Alonso, D., Fernandez-Eslava, B. and Alonso-Alvarez, C. (2020a), Testing the shared-pathway hypothesis in the carotenoid-based coloration of red crossbills. Evolution 74, 2348–2364.

Cantarero, A., Andrade, P., Carneiro, M., Moreno-Borrallo, A. and Alonso-Alvarez, C. (2020b). Testing the carotenoid-based sexual signalling mechanism by altering CYP2J19 gene expression and colour in a bird species. Proceedings of the Royal Society B 287, 20201067.

Casagrande, S., and Hau, M. (2018). Enzymatic antioxidants but not baseline glucocorticoids mediate the reproduction–survival trade-off in a wild bird. Proceedings of the Royal Society B 285, doi:10.1098/rspb.2018.2141

Chui, C. K. S., McGraw, K. J. and Doucet, S. M. (2011). Carotenoid-based plumage coloration in golden-crowned kinglets Regulus satrapa: Pigment characterization and relationships with migratory timing and condition. Journal of Avian Biology 42, 309–322. doi:10.1111/j.1600-048X.2011.05240.x

Cornelius, J. M. and Hahn, T. P. (2012). Seasonal pre-migratory fattening and increased activity in a nomadic and irruptive migrant, the Red Crossbill Loxia curvirostra. Ibis 154, 693–702. doi:10.1111/j.1474-919X.2012.01266.x

Costantini, D., Dell’Ariccia, G. and Lipp, H. P. (2008). Long flights and age affect oxidative status of homing pigeons (Columba livia). Journal of Experimental Biology 211, 377–381. doi:10.1242/jeb.012856

Dawson, A. (2015) Avian Molting, in Pages 907-917, Sturkie’s Avian Physiology, Chapter 38 (Sixth Edition). Editor(s): Colin G. Scanes. London UK: Academic Press.

Dawson, W. R. and Tordoff, H. B. (1964). Relation of Oxygen Consumption to Temperature in the Red and White-Winged Crossbills. Auk 81, 26–35. doi:10.2307/4082607

Del Val, E., Senar, J. C., Garrido-Fernández, J., Jarén, M., Borràs, A., Cabrera, J. and Negro, J. J. (2009a). The liver but not the skin is the site for conversion of a red carotenoid in a passerine bird. Naturwissenschaften 96, 797–801. doi:10.1007/s00114-009-0534-9

Del Val, E., Borràs, A., Cabrera, J., Senar, J. C. and Quesada, J. (2009b). Plumage colour of male Common Crossbills Loxia curvirostra⍰: visual assessment validated by colorimetry. Revista Catalana d ‘Ornitologia 25, 19–25.

Deviche, P., Mcgraw, K. J. and Underwood, J. (2008). Season-, sex-, and age-specific accumulation of plasma carotenoid pigments in free-ranging white-winged crossbills Loxia leucoptera. Journal of Avian Biology 39, 283–292. doi:10.1111/j.2008.0908-8857.04164.x

Draud, T. (2019). The cost of breeding in the winter versus the summer in an opportunistic, northtemperate songbird, the red crossbill (Loxia curvirostra). PhD Thesis. Michigan, USA: Eastern Michigan University.

Fernández-Eslava, B., Alonso, D. and Alonso-Alvarez, C. (2021a). An age-related decline in the expression of a red carotenoid-based ornament in wild birds. Evolution 75, 3142–3153. doi:10.1111/evo.14378

Fernández-Eslava, B., Alonso, D., Galicia, D. and Arizaga, J. (2021b). Strong evidence supporting a relationship between colour pattern and apparent survival in common crossbills. Journal of Ornithology 163, 243–249. doi:10.1007/s10336-021-01927-4

Fernández-Eslava, B., Alonso, D., Galicia, D. and Arizaga, J. (2020). Estimation of moult duration in birds with suspended moults: the case of the Red Crossbill and its relation to reproduction. Journal of Ornithology 161, 481–490. doi:10.1007/s10336-019-01739-7

Fernández-Eslava, B., Cantarero, A., Alonso, D. and Alonso-Alvarez, C. Data set of “Wild common crossbills produce redder body feathers when their wings are clipped” article. DIGITAL.CSIC, DOI and link upon acceptance.

Foley, J.D. and Van Dam, A. (1982). Fundamentals of interactive computer graphics, Reading, MA: Addison-Wesley,

Galván, I., Murtada, K., Jorge, A., Riós, Á., and Zougagh, M. (2019). Unique evolution of Vitamin A as an external pigment in tropical starlings. Journal of Experimental Biology, 222(11), 1–7. doi:10.1242/jeb.205229

Gerson, A. R. (2012). Environmental Physiology of Flight in Migratory Birds. Electronic Thesis and Dissertation Repository. 734. Ontario, Canada: University of Western Ontario. https://ir.lib.uwo.ca/etd/734

Gómez, J. and Liñán-Cembrano, G. (2017). SpotEgg: an image-processing tool for automatised analysis of colouration and spottiness. Journal of Avian Biology 48, 502–512.

Grafen, A. (1990). Biological signals as handicaps. J. Theor. Biol. 144, 517–546.

Gulde, V. A. L., Renema, R. and Bédécarrats, G. Y. (2010). Use of dietary thyroxine as an alternate molting procedure in spent turkey breeder hens. Poultry Science 89, 96–107. doi:10.3382/ps.2009-00294

Gutiérrez, J. S., Sabat, P., Castañeda, L. E., Contreras, C., Navarrete, L., Peña-Villalobos, I. and Navedo, J. G. (2019). Oxidative status and metabolic profile in a long-lived bird preparing for extreme endurance migration. Scientific Reports 9, 1–11. doi:10.1038/s41598-019-54057-6

Hahn, T. P. (1995). Integration of photoperiodic and food cues to time changes in reproductive physiology by an opportunistic breeder, the red crossbill, Loxia curvirostra (Aves: Carduelinae). Journal of Experimental Zoology 272, 213–226. doi:10.1002/jez.1402720306

Harding, A. M. A., Kitaysky, A. S., Hamer, K. C., Hall, M. E., Welcker, J., Talbot, S. L., Karnovsky N.J. Gabrielsen G.W. and Grémillet, D. (2009). Impacts of experimentally increased foraging effort on the family: offspring sex matters. Animal Behaviour 78, 321–328. doi:10.1016/j.anbehav.2009.05.009

Hill, G. E. (2011). Condition-dependent traits as signals of the functionality of vital cellular processes. Ecology Letters 14, 625–634.

Hill, G. E., Hood, W. R., Ge, Z., Grinter, R., Greening, C., James, D., Park, N. R., Taylor, H. A., Andreasen, V. A., Powers, M. J. et al. (2019). Plumage redness signals mitochondrial function in the House Finch. Proc. R. Soc. B 286, 20191354.

Jenni-Eiermann, S., Jenni, L., Smith, S., and Costantini, D. (2014). Oxidative stress in endurance flight: An unconsidered factor in bird migration. PLoS ONE 9, 1–6. doi:10.1371/journal.pone.0097650

Johnson, J. D., and Hill, G. E. (2013). Is carotenoid ornamentation linked to the inner mitochondria membrane potential? A hypothesis for the maintenance of signal honesty. Biochimie 95, 436–444. doi:10.1016/j.biochi.2012.10.021

Jones, I. L. (1994) Mass changes of Least Auklets Aethia pusilla during the breeding season: evidence for programmed loss of mass. Journal of Animal Ecology 63, 71–78.

Kaiser, J. (1993). A new multi-category classification of subcutaneous fat deposits of songbirds. Journal of Field Ornithology 64, 246–255.

Koch, R. E., Kavazis, A. N., Hasselquist, D., Hood, W. R., Zhang, Y., Toomey, M. B. and Hill, G. E. (2018). No evidence that carotenoid pigments boost either immune or antioxidant defenses in a songbird. Nature Communications, 9, 491. doi:10.1038/s41467-018-02974-x

Leclaire, S., Bourret, V., Pineaux, M., Blanchard, P., Danchin, E., and Hatch, S. A. (2019). Red coloration varies with dietary carotenoid access and nutritional condition in kittiwakes. Journal of Experimental Biology 222, jeb210237. doi:10.1242/jeb.210237

Leclaire, S., Bourret, V., Wagner, R. H., Hatch, S. A., Helfenstein, F., Chastel, O. and Danchin, É. (2011). Behavioral and physiological responses to male handicap in chick-rearing black-legged kittiwakes. Behavioral Ecology 22, 1156–1165. doi:10.1093/beheco/arr149

Lessells, C. M. and Boag, P. T. (1987). Unrepeatable repeatabilities: a common mistake. Auk 104, 116–121.

Lifjeld, J. T. and Slagsvold, T. (1988). Female pied flycatchers Ficedula hypoleuca choose male characteristics in homogeneous habitats. Behavioral Ecology and Sociobiology 22, 27–36. doi:10.1007/BF00395695

Lind, J. and Jakobsson, S. (2001). Body building and concurrent mass loss: Flight adaptations in tree sparrows. Proceedings of the Royal Society B 268, 1915–1919. doi:10.1098/rspb.2001.1740

Lopes, R. J. J., Johnson, J. D. D., Toomey, M. B. B., Ferreira, M. S. S., Araujo, P. M. M., Melo-Ferreira, J. Andersson, L., Hill, G. E., Corbo, J. C., and Carneiro, M. (2016). Genetic Basis for Red Coloration in Birds. Current Biology 26, 1427–1434. doi:10.1016/j.cub.2016.03.076

Mateos-Gonzalez, F., Hill, G., and Hood, W. (2014). Carotenoid coloration predicts escape performance in the House Finch (Haemorhous mexicanus). Auk 131, 275–281. doi:10.1642/AUK-13-207.1

Matysioková, B., and Remeš, V. (2011). Responses to increased costs of activity during incubation in a songbird with female-only incubation: Does feather colour signal coping ability? Journal of Ornithology 152, 337–346. doi:10.1007/s10336-010-0594-9

Maynard Smith, J. and D. Harper. (2003). Animal signals. Oxford Series in Ecology and Evolution. Oxford, UK: Oxford University Press-

McGraw, K. J. 2006. Mechanics of carotenoid-based coloration. in G. E. Hill and K. J. McGraw, eds. Bird coloration, Volume 1: Mechanisms and measurements. Harvard: Harvard University Press.

McGraw, K. J. and Toomey, M. B. (2010). Carotenoid accumulation in the tissues of zebra finches: predictors of integumentary pigmentation and implications for carotenoid allocation strategies. Physiological and Biochemical Zoology 83, 97–109. doi:10.1086/648396

Merkle, M. S. and Barclay, R. M. R. (1996). Body Mass Variation in Breeding Mountain Bluebirds Sialia currucoides: Evidence of Stress or Adaptation for Flight? Journal of Animal Ecology 65, 401. doi:10.2307/5776

Moreno, J. (1989). Energetic constraints on uniparental incubation in the wheatear Oenanthe oenanthe (L). Ardea 77, 107e115.

Park, S. Y., Rossman, M. J., Gifford, J. R., Bharath, L. P., Bauersachs, J., Richardson, R. S., E. Abel D., Symons, D. and Riehle, C. (2016). Exercise training improves vascular mitochondrial function. American Journal of Physiology - Heart and Circulatory Physiology 310, H821–H829. doi:10.1152/ajpheart.00751.2015

Penn, D. J. and S. Számadó. 2020. The Handicap Principle: how an erroneous hypothesis became a scientific principle. Biological Reviews 95, 267–290.

Pérez, J. H., Meddle, S. L., Wingfield, J. C. and Ramenofsky, M. (2018). Effects of thyroid hormone manipulation on pre-nuptial molt, luteinizing hormone and testicular growth in male white-crowned sparrows (Zonotrichia leuchophrys gambelii). General and Comparative Endocrinology 255, 12–18. doi:10.1016/j.ygcen.2017.09.025

Potti, J. (1998). Arrival time from spring migration in male pied flycatchers: individual consistency and familial resemblance. Condor 100, 702–708. doi:10.2307/1369752

Rowe, M. and McGraw, K. J. (2008). Carotenoids in the seminal fluid of wild birds: Interspecific variation in fairy-wrens. Condor 110(4), 694–700. doi:10.1525/cond.2008.8604

Salin, K., Auer, S. K., Rey, B., Selman, C. and Metcalfe, N. B. (2015). Variation in the link between oxygen consumption and ATP production, and its relevance for animal performance. Proceedings of the Royal Society B 282, 20151028. doi:10.1098/rspb.2015.1028

Schmidt-Wellenburg, C. A., Visser, G. H., Biebach, B., Delhey, K., Oltrogge, M., Wittenzellner, A., Biebach., H. and Kempenaers, B. (2008). Trade-off between migration and reproduction: Does a high workload affect body condition and reproductive state? Behavioral Ecology 19, 1351–1360. doi:10.1093/beheco/arn066

Schut, E., Magrath, M. J. L., Van Oers, K. and Komdeur, J. (2012). Volume of the cloacal protuberance as an indication of reproductive state in male Blue Tits Cyanistes caeruleus. Ardea, 100, 202–205. doi:10.5253/078.100.0212

Sies H, Jones DP. (2007). Oxidative stress. in: ‘Encyclopedia of Stress’ (Fink, G., ed.), 2nd Ed., Vol. 3, Pp 43–49. Amsterdam, Netherlands: Elsevier.

Simons, M. J. P., Cohen, A. A. and Verhulst, S. (2012). What does carotenoid-dependent coloration tell? Plasma carotenoid level signals immunocompetence and oxidative stress state in birds-A meta-analysis. PloS One 7, e43088. doi:10.1371/journal.pone.0043088

Slagsvold, T. and Johansen, M. (1998). Mass loss in female pied flycatchers Ficedula hypoleuca during late incubation: Supplementation fails to support the reproductive stress hypothesis. Ardea 86, 203–211.

Slagsvold, T. and Lifjeld, J. T. (1990). Influence of male and female quality on clutch size in tits (Parus Spp.) Ecology 71, 1258–1266.

St-Pierre, J., Buckingham, J. A., Roebuck, S. J. and Brand, M. D. (2002). Topology of superoxide production from different sites in the mitochondrial electron transport chain. Journal of Biological Chemistry 277, 44784–44790. doi:10.1074/jbc.M207217200

Tarvin, K. A., Wong, L. J., Lumpkin, D. C., Schroeder, G. M., D’Andrea, D., Meade, S., Rivers, P. and Murphy, T. G. (2016). Dynamic status signal reflects outcome of social interactions, but not energetic stress. Frontiers in Ecology and Evolution 4, 1–12. doi:10.3389/fevo.2016.00079

Teulier, L., Rouanet, J. L., Letexier, D., Romestaing, C., Belouze, M., Rey, B., Duchamp, C. and Roussel, D. (2010). Cold-acclimation-induced non-shivering thermogenesis in birds is associated with upregulation of avian UCP but not with innate uncoupling or altered ATP efficiency. Journal of Experimental Biology 213, 2476–2482. doi:10.1242/jeb.043489

Tonra, C. M., Marini, K. L. D., Marra, P. P., Germain, R. R., Holberton, R. L. and Reudink, M. W. (2014). Color expression in experimentally regrown feathers of an overwintering migratory bird: Implications for signaling and seasonal interactions. Ecology and Evolution 4, 1222–1232. doi:10.1002/ece3.994

Völker, V. O. (1957). Die experimentelle Rotfärbung des Gefieders beim Fichtenkreuzschnabel (Loxia curvirostra). Journal Fur Ornithologie 98, 210–214.

Walter, I. and Seebacher, F. (2009). Endothermy in birds: Underlying molecular mechanisms. Journal of Experimental Biology 212, 2328–2336. doi:10.1242/jeb.029009

Weber, H. (1953). Bewirkung des Farbwechsels bei männlichen Kreuzschnäbeln. Journal Für Ornithologie 94, 342–346. doi:10.1007/BF01922518

Weber, V. H. (1961). Uber die Ursache des Verlustes der roten Federfarbe bei gek ∼ ifigten Birkenzeisigen Von Hubert Weber Die experimentelle Priifung. Journal Fur Ornithologie 102, 158–163.

Wegmann, M., Voegeli, B. and Richner, H. (2015). Oxidative status and reproductive effort of great tits in a handicapping experiment. Behavioral Ecology 26, 747–754. doi:10.1093/beheco/arv006

Weimerskirch, H., Chastel, O. and Ackermann, L. (1995). Adjustment of parental effort to manipulated foraging ability in a pelagic seabird, the thin-billed prion Pachyptila belcheri. Behavioral Ecology and Sociobiology 36, 11–16. doi:10.1007/BF00175723

Yap, K. N., Kim, O. R., Harris, K. C. and Williams, T. D. (2017). Physiological effects of increased foraging effort in a small passerine. Journal of Experimental Biology 220, 4282–4291. doi:10.1242/jeb.160812

Zahavi, A. (1975). Mate selection a selection for a handicap. Journal of Theoretical Biology 53, 205–214.

Zhang, Y. and Wong, H. S. (2021). Are mitochondria the main contributor of reactive oxygen species in cells? Journal of Experimental Biology 224, jeb221606 doi:10.1242/jeb.221606

